# Whole genome and transcriptome maps of the entirely black native Korean chicken breed *Yeonsan Ogye*

**DOI:** 10.1101/224311

**Authors:** Jang-il Sohn, Kyoungwoo Nam, Hyosun Hong, Jun-Mo Kim, Dajeong Lim, Kyung-Tai Lee, Yoon Jung Do, Chang Yeon Cho, Namshin Kim, Han-Ha Chae, Jin-Wu Nam

## Abstract

*Yeonsan Ogye (YO)*, an indigenous Korean chicken breed (*gallus gallus domesticus*), has entirely black external features and internal organs. In this study, the draft genome of *YO* was assembled using a hybrid *de novo* assembly method that takes advantage of high-depth Illumina short-reads (232.2X) and low-depth PacBio long-reads (11.5X). Although the contig and scaffold N50s (defined as the shortest contig or scaffold length at 50% of the entire assembly) of the initial *de novo* assembly were 53.6Kbp and 10.7Mbp, respectively, additional and pseudo-reference-assisted assemblies extended the assembly to 504.8Kbp for contig N50 (pseudo-contig) and 21.2Mbp for scaffold N50, which included 551 structural variations including the Fibromelanosis (*FM*) locus duplication, compared to galGal4 and 5. The completeness (97.6%) of the draft genome (Ogye_1) was evaluated with single copy orthologous genes using BUSCO, and found to be comparable to the current chicken reference genome (galGal5; 97.4%), which was assembled with a long read-only method, and superior to other avian genomes (92~93%), assembled with short read-only and hybrid methods. To comprehensively reconstruct transcriptome maps, RNA sequencing (RNA-seq) and representation bisulfite sequencing (RRBS) data were analyzed from twenty different tissues, including black tissues. The maps included 15,766 protein-coding and 6,900 long non-coding RNA genes, many of which were expressed in the tissue-specific manner, closely related with the DNA methylation pattern in the promoter regions.

## Background

The *Yeonsan Ogye* (*YO*), a designated natural monument of Korea (No. 265), is an indigenous Korean chicken breed that is notable for its entirely black plumage, skin, beak, comb, eyes, shank, claws, and internal organs [1]. In terms of its plumage and body color, as well as its number of toes, this unique chicken breed resembles the indigenous Indonesian chicken breed *Ayam cemani* [2-4]. *YO* also has some morphological features that are similar to those of the *Silkie* fowl, except for a veiled black walnut comb and hair-like, fluffy plumage that is white or variably colored [5, 6]. Although the exact origin of the *YO* breed has not yet been clearly defined, its features and medicinal usages were recorded in *Dongui Bogam* [7], a traditional Korean medical encyclopedia compiled and edited by Heo Jun in 1613.

To date, a number of avian genomes from both domestic and wild species have been constructed and compared, revealing genomic signatures associated with the domestication process and genomic differences that provide an evolutionary perspective [8]. The chicken reference genome was first assembled using the *Red junglefowl* [9], first domesticated at least five thousand years ago in Asia; the latest version of the reference genome was released in 2015 (galGal5, GenBank Assembly ID GCA_000002315.3) [10]. However, because domesticated chickens exhibit diverse morphological features, including skin and plumage colors, the genome sequences of unique breeds are necessary for understanding their characteristic phenotypes through analyses of single nucleotide polymorphisms (SNPs), insertions and deletions (INDELs), structural variations (SVs), and coding and non-coding transcriptomes. Here, we provide the first version of *YO* genome (Ogye_1) and transcriptome maps, which include annotations of large SVs, SNPs, INDELs, and repeats, as well as coding and non-coding transcriptome maps along with DNA methylation landscapes across twenty different tissues of *YO.*

## Results

### Sample collection and data description

8-month-old *YO* chickens (object number: 02127), obtained from the Animal Genetic Resource Research Center of the National Institute of Animal Science (Namwon, Korea), were used in the study (**Figure 1A**). The protocols for the care and experimental use of *YO* were reviewed and approved by the Institutional Animal Care and Use Committee of the National Institute of Animal Science (IACUC No.: 2014-080). *YO* management, treatment, and sample collection took place at the National Institute of Animal Science.

**Figure 1.**
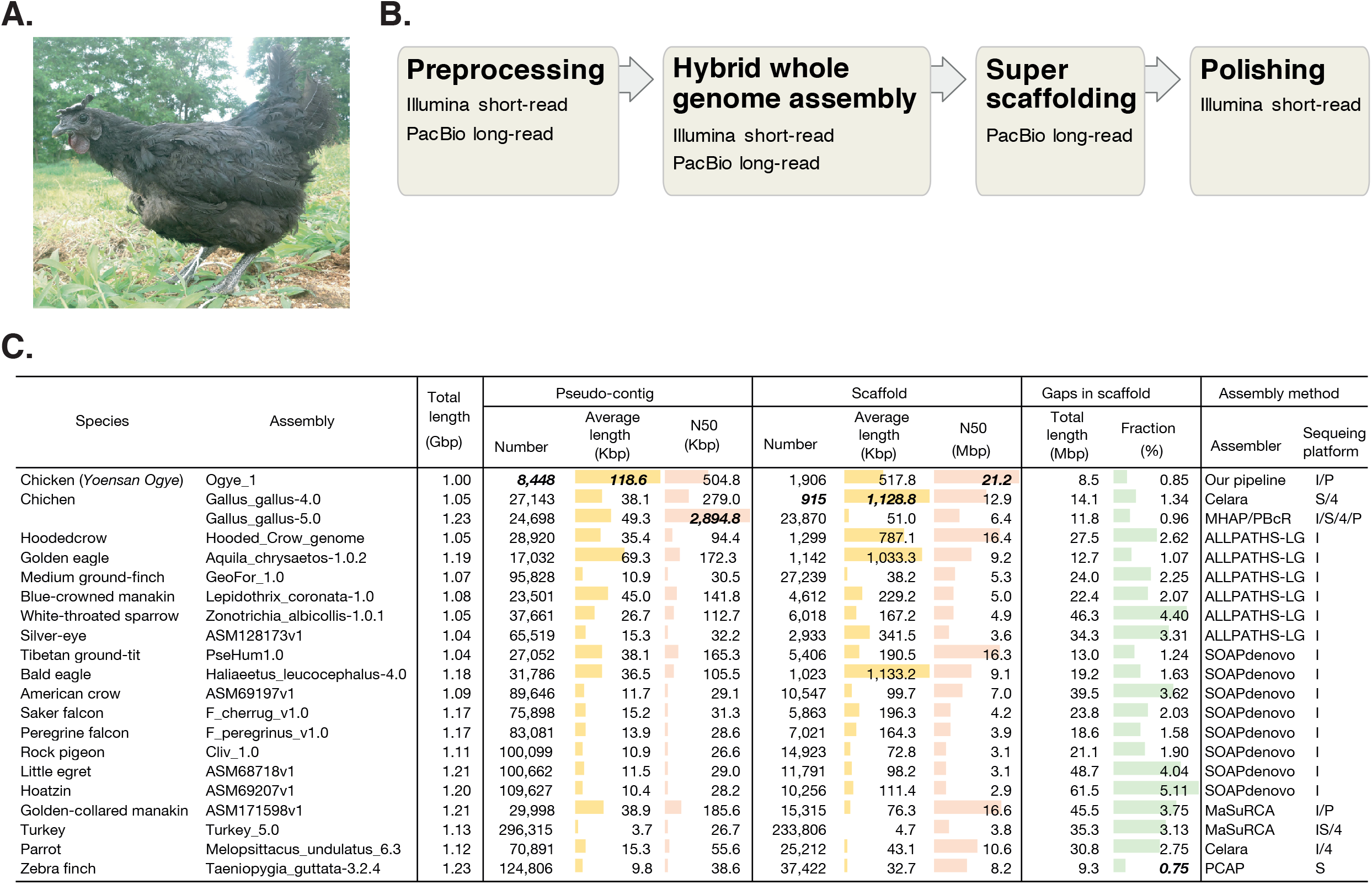
**A.** Picture of *Yeonsan Ogye*; **B.** Hybrid genome assembly pipeline; **C.** The N50 and average length of pseudo-contigs and scaffolds of the Ogye_1 and other avian genomes created using the indicated assembly methods (last column; here, sequencing platforms are designated as follows: “I” indicates Illumina, “P” is PacBio, “S” is Sanger, and “4” is Roche454).

#### Whole genome sequencing

Genomic DNA was extracted from blood using Wizard DNA extraction kit [11] and prepared for DNA sequencing library construction. According to the DNA fragment (insert) size, three different library types were constructed: paired-end library for small inserts (280 and 500 bp) and mate-pair library for large inserts (3, 5, 8, and 10 Kbp), and fosmid libraries (40 Kbp) using Illumina’s protocols (Illumina, San Diego, CA, USA) (**Table 1**). The constructed libraries were sequenced using Illumina’s Hiseq2000 platform. In total, 232.2 X Illumina short reads were obtained (59.6 X from the small insert libraries and 172.6 X from the large insert libraries) and, after filtering raw data with low quality (> 30% of the base-pairs in a read have a Phred score <20), 163.5X were used for genome assembly. To fill gaps and improve the scaffold quality, 11.5X PacBio long reads were additionally sequenced; the average length of the long reads was 6Kbp (**Table 1**).

**Table 1.**
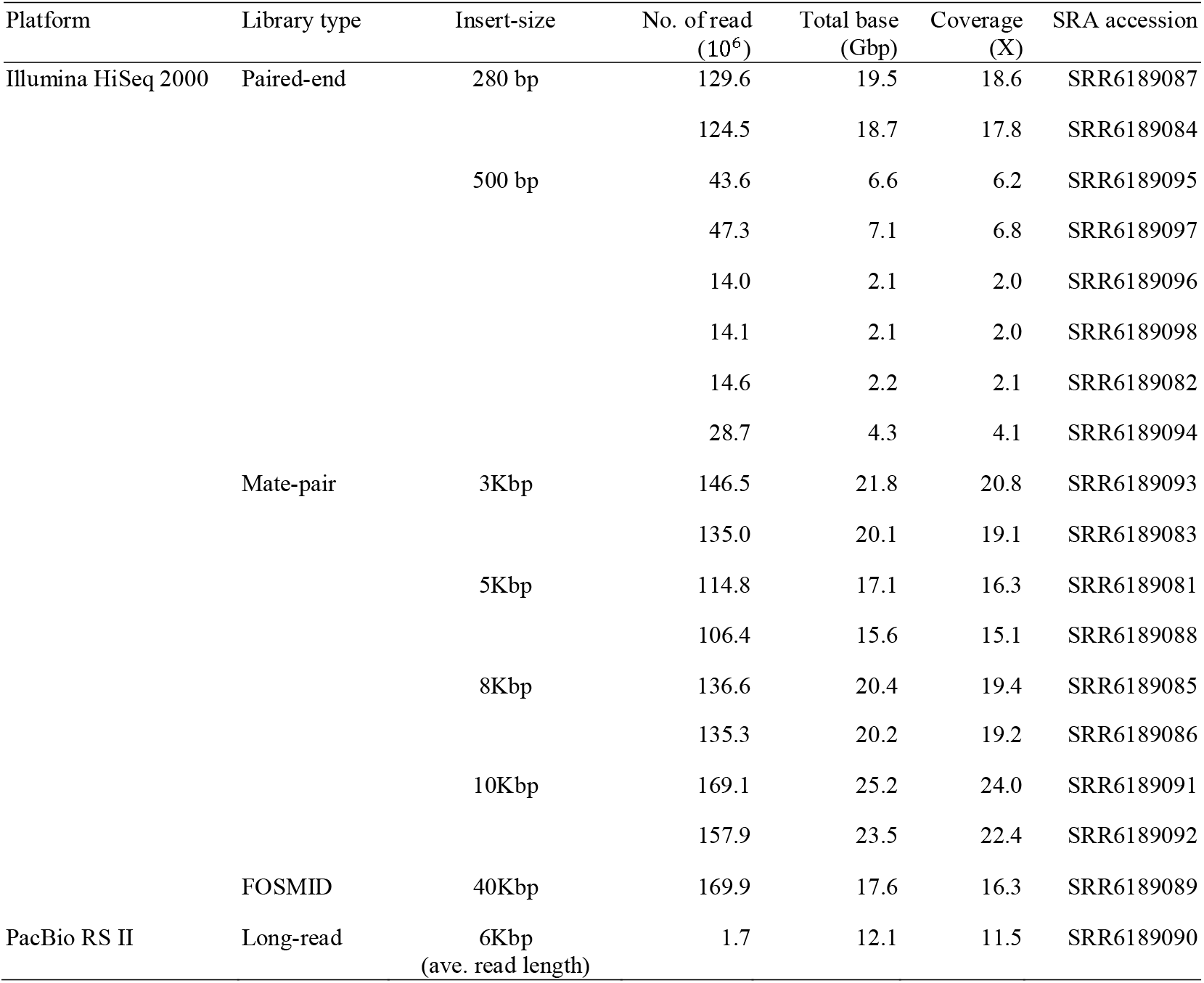
Whole genome sequencing data.

#### Whole transcriptome sequencing

Total RNAs were extracted from twenty different tissues using 80% EtOH and TRIzol. The RNA concentration was checked by Quant-IT RiboGreen (Invitrogen, Carlsbad, USA). To assess the integrity of the total RNA, samples were run on a TapeStation RNA screentape (Agilent, Waldbronn, Germany). Only high quality RNA samples (RIN ≥R7.0) were used for RNA-seq library construction. Each library was independently prepared with 300ng of total RNA using an Illumina TruSeq Stranded Total RNA Sample Prep Kit (Illumina, San Diego, CA, USA). The rRNA in the total RNA was depleted using a Ribo-Zero kit. After rRNA depletion, the remaining RNA was purified, fragmented and primed for cDNA synthesis. The cleaved RNA fragments were copied into the first cDNA strand using reverse transcriptase and random hexamers. This step was followed by second strand cDNA synthesis using DNA Polymerase I, RNase H and dUTP. The resulting cDNA fragments then underwent an end repair process, the addition of a single ‘A’ base, after which adapters were ligated. The products were purified and enriched with PCR to create the final cDNA library. The libraries were quantified using qPCR according to the qPCR Quantification Protocol Guide (KAPA Library Quantificatoin kits for Illumina Sequecing platforms) and qualified using the TapeStation D1000 ScreenTape assay (Agilent Technologies, Waldbronn, Germany). As a result, we have sequenced about 1.5 billion RNA-seq reads from twenty different tissues, which are Breast, Liver, Bone marrow, Fascia, Cerebrum, Gizzard, Immature egg, Comb, Spleen, Mature egg, Cerebellum, Gall bladder, Kidney, Heart, Uterus, Pancreas, Lung, Skin, Eye, and Shank (**Table 2**).

**Table 2.** Sequencing and mapping summary of RNA-seq reads.

| Samples | Paired-end |  |  | Single-end |  |  |
| --- | --- | --- | --- | --- | --- | --- |
|  | No. of reads | Mapping rate | SRA accession | No. of reads | Mapping rate | SRA accession |
| Breast | 34,893,064 | 92.05% | SRX3223583 | 43,294,022 | 90.70% | SRX3223603 |
| Liver | 33,476,266 | 85.75% | SRX3223584 | 48,032,813 | 85.81% | SRX3223604 |
| Bone marrow | 30,975,506 | 85.00% | SRX3223585 | 40,286,974 | 87.99% | SRX3223605 |
| Fascia | 33,316,764 | 84.61% | SRX3223586 | 42,425,452 | 87.93% | SRX3223606 |
| Cerebrum | 30,887,821 | 89.95% | SRX3223587 | 46,455,658 | 92.32% | SRX3223607 |
| Gizzard | 31,537,118 | 84.00% | SRX3223588 | 38,689,871 | 85.82% | SRX3223608 |
| Immature egg | 32,009,437 | 87.73% | SRX3223589 | 32,048,703 | 87.80% | SRX3223609 |
| Comb | 31,936,332 | 85.34% | SRX3223590 | 37,985,049 | 87.76% | SRX3223610 |
| Spleen | 28,946,777 | 89.70% | SRX3223591 | 38,704,448 | 89.33% | SRX3223611 |
| Mature egg | 30,873,699 | 91.98% | SRX3223592 | 40,650,664 | 92.17% | SRX3223612 |
| Cerebellum | 30,798,145 | 93.53% | SRX3223593 | 39,940,946 | 93.34% | SRX3223613 |
| Gall bladder | 35,862,229 | 84.83% | SRX3223594 | 35,423,339 | 87.06% | SRX3223614 |
| Kidney | 29,953,007 | 87.25% | SRX3223595 | 39,894,009 | 89.99% | SRX3223615 |
| Heart | 30,986,431 | 94.14% | SRX3223596 | 45,951,338 | 91.49% | SRX3223616 |
| Uterus | 33,444,002 | 91.89% | SRX3223597 | 46,650,355 | 90.63% | SRX3223617 |
| Pancreas | 30,595,568 | 82.52% | SRX3223598 | 47,361,192 | 84.35% | SRX3223618 |
| Lung | 31,533,498 | 87.63% | SRX3223599 | 45,552,982 | 92.34% | SRX3223619 |
| Skin | 34,442,464 | 82.36% | SRX3223600 | 41,934,970 | 84.00% | SRX3223620 |
| Eye | 33,006,509 | 89.21% | SRX3223601 | 44,044,630 | 91.82% | SRX3223621 |
| Shank | 28,643,334 | 94.07% | SRX3223602 | 47,716,995 | 79.86% | SRX3223622 |

#### Reduced representation bisulfìte sequencing

Preparation of reduced representation bisulfite sequencing (RRBS) libraries was done following Illumina’s RRBS protocol. 5μg of genomic DNA that had been digested with the restriction enzyme MspI and purified with a QIAquick PCR purification kit (QIAGEN, Hilden, Germany) was used for library preparation, which was done using a TruSeq Nano DNA Library Prep Kit (Illumina, San Diego, USA). Eluted DNA fragments were end-repaired, extended on the 3’ end with an ‘A’, and ligated with Truseq adapters. After ligation had been assessed, the products, which ranged from 175 to 225bp in length (insert DNA of 55–105 bp plus adaptors of 120 bp), were excised from a 2%(w/v) Low Range Ultra Agarose gel (Biorad, Hercules, USA) and purified using the QIAquick gel extraction protocol. The purified DNA underwent bisulfite conversion using an EpiTect Bisulfite Kit (Qiagen, 59104). The bisulfite-converted DNA libraries were amplified by PCR (four cycles) using PfuTurbo Cx DNA polymerase (Agilent, 600410). The final product was then quantified using qPCR and qualified using the Agilent Technologies 2200 TapeStation assay (Agilent, Waldbronn, Germany). The final product was sequenced using the HiSeq™ 2500 platform (Illumina, San Diego, USA). As a result, we have produced 123 million RRBS reads (see **Table 3**) from twenty different tissues.

**Table 3.** Sequencing and mapping summary of RRBS reads.

| Samples | No. of reads | Mapping rate | SRA accession |
| --- | --- | --- | --- |
| Breast | 6,042,106 | 68.90% | SRX3223667 |
| Liver | 6,744,208 | 74.20% | SRX3223668 |
| Bone marrow | 5,736,011 | 72.00% | SRX3223669 |
| Fascia | 5,720,194 | 68.90% | SRX3223670 |
| Cerebrum | 6,078,989 | 70.00% | SRX3223671 |
| Gizzard | 5,731,878 | 69.40% | SRX3223672 |
| Immature egg | 6,741,258 | 67.70% | SRX3223673 |
| Comb | 5,948,687 | 72.90% | SRX3223674 |
| Spleen | 6,307,517 | 77.60% | SRX3223675 |
| Mature egg | 6,246,607 | 69.20% | SRX3223676 |
| Cerebellum | 6,291,610 | 68.20% | SRX3223677 |
| Gall bladder | 5,738,180 | 70.10% | SRX3223678 |
| Kidney | 5,470,502 | 68.60% | SRX3223679 |
| Heart | 5,462,739 | 69.40% | SRX3223680 |
| Uterus | 6,046,764 | 67.90% | SRX3223681 |
| Pancreas | 7,100,215 | 70.30% | SRX3223682 |
| Lung | 5,640,120 | 67.60% | SRX3223683 |
| Skin | 7,226,309 | 72.40% | SRX3223684 |
| Eye | 6,956,141 | 71.90% | SRX3223685 |
| Shank | 5,924,463 | 74.20% | SRX3223686 |

### Hybrid whole genome assembly

The Ogye_1 genome was assembled using our hybrid genome assembly pipeline, employing the following four steps: preprocessing, hybrid *de novo* assembly, super-scaffolding, and polishing (**Figure 1B** and **Figure S1**). During the preprocessing step, the errors in the Illumina short-reads were corrected by KmerFreq and Corrector [12] using sequencing quality scores. In turn, using the corrected short-reads, the sequencing errors in the PacBio long reads were corrected by LoRDEC [13].

In hybrid *de novo* genome assembly, the initial assembly was done with the error-corrected short reads from the paired-end and mate-pair libraries using ALLPATHS-LG [14] with the default option, producing contigs and scaffolds. The resulting contigs and scaffolds showed 53.6 Kbp and 10.7 Mb of N50 (**Figure S1**), respectively. Next, the scaffolds were additionally connected using SSPACE-LongRead [15] and OPERA [16] with corrected PacBio long reads and FOSMID reads. The gaps within and between scaffolds were re-examined with GapCloser [12] with error-corrected short-reads. All resulting scaffolds were aligned to the galGal4 genome (GenBank assembly accession: GCA_000002315.2) by LASTZ [17] to detect putative mis-assemblies, verified by paired-end and mate-pair reads mapped to the scaffolds using BWA-MEM [18]. Comparison with results of LASTZ and BWA-MEM detected 30 misassemblies, break points of which were detected (**Figure S2.**) using Integrative Genomics Viewer (IGV) [19] and in-house programs. Breaking scaffolds at the break points resulted in pseudo-contig N50 of 108.6 Kbp and scaffold N50 of 18.7 Mb (**Figure S1**). A pseudo-contig is defined as a sequence broken by gaps of >1bp or a single N, which are assumed to be gaps or errors.

In the super-scaffolding stage, pseudo-reference-assisted assembly was done using LASTZ, BWA-MEM, PBJelly [20], and SSPACE-LongRead to enhance the quality of assembly using error-corrected PacBio long-reads to reduce the topological complexity of the assembly graphs [21]. Because even scaffolding with long-reads can be affected by repetitive sequences, the results of mapping scaffolds to each chromosome were transformed into a hierarchical bipartite graph (**Figure S3**) to minimize the influence of repetitive sequences. The hierarchical bipartite graph was built by mapping PacBio (error-corrected) reads to scaffolds using BWA-MEM and, in turn, mapping scaffolds to the galGal4 genome (GCA_000002315.2) using LASTZ. Using the hierarchical bipartite graphs, all scaffolds and PacBio reads were finally assigned to each chromosome. Based on the results, super-scaffolding and additional gap-filling was performed by SSPACE-LongRead and PBJelly, respectively, resulting in scaffold N50 of 21.2Mbp (**Figure 1C** and **Figure S1**). In the last stage, nucleotide errors or ambiguities were corrected by the GATK pipeline [22] with paired-end reads, and in turn, any vector contamination was removed using VecScreen with UniVec database [23]. The final assembly results showed that the gap percentage and (pseudo-)contig N50 were significantly improved, from 1.87% and 53.6 Kbp in the initial assembly to 0.85% and 504.8 Kbp in the final assembly, respectively. Among avian genome assemblies, these results are second best and the scaffold N50 is the best (**Figure 1C**). The complete genome sequence at the chromosome level (**Figure S4**) was built by connecting final scaffolds in the order of appearance in each chromosome with the introduction of 100 Kbp ‘N’ gaps between them. To evaluate the completeness of the genome, the *YO* draft genome was compared to the galGal4 (short read-based assembly) and galGal5 (long read-based assembly) genomes, with respect to 2,586 conserved vertebrate genes, using BUSCO [24]. The results indicated that the Ogye_1 genome contains more complete single-copy BUSCO genes, suggesting that the Ogye_1 genome is slightly more complete than the others (**Table 4**).

**Table 4.**
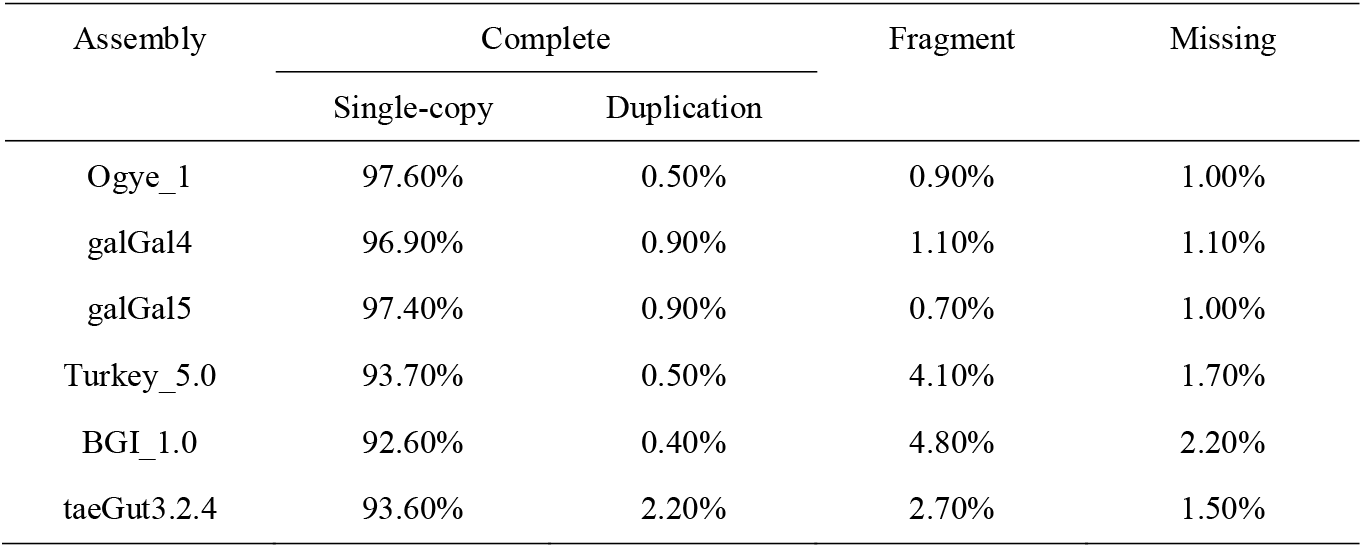
Comparison of genome completeness using BUSCO.

### Large structural variations

When the Ogye_1 genome assembly was compared to two versions of the chicken reference genome assembly, galGal4 and 5, using LASTZ [17] and in-house programs/scripts, 551 common large (≥1 Kbp) structural variations (SVs) evident in both assembly versions were detected by at least two different SV prediction programs (Delly, Lumpy, FermiKit, and novoBreak) [25-28] (**Figure 1A**; **Table S1**). SVs included 185 deletions (DELs), 180 insertions (INSs), 158 duplications (DUPs), 23 inversions (INVs), and 5 intra or inter-chromosomal translocations (TRs). 290 and 447 distinct SVs were detected relative to galGal 4 and galGal5, respectively (**Figure 2A**), suggesting that the two reference assemblies could include mis-assembly.

**Figure 2.**
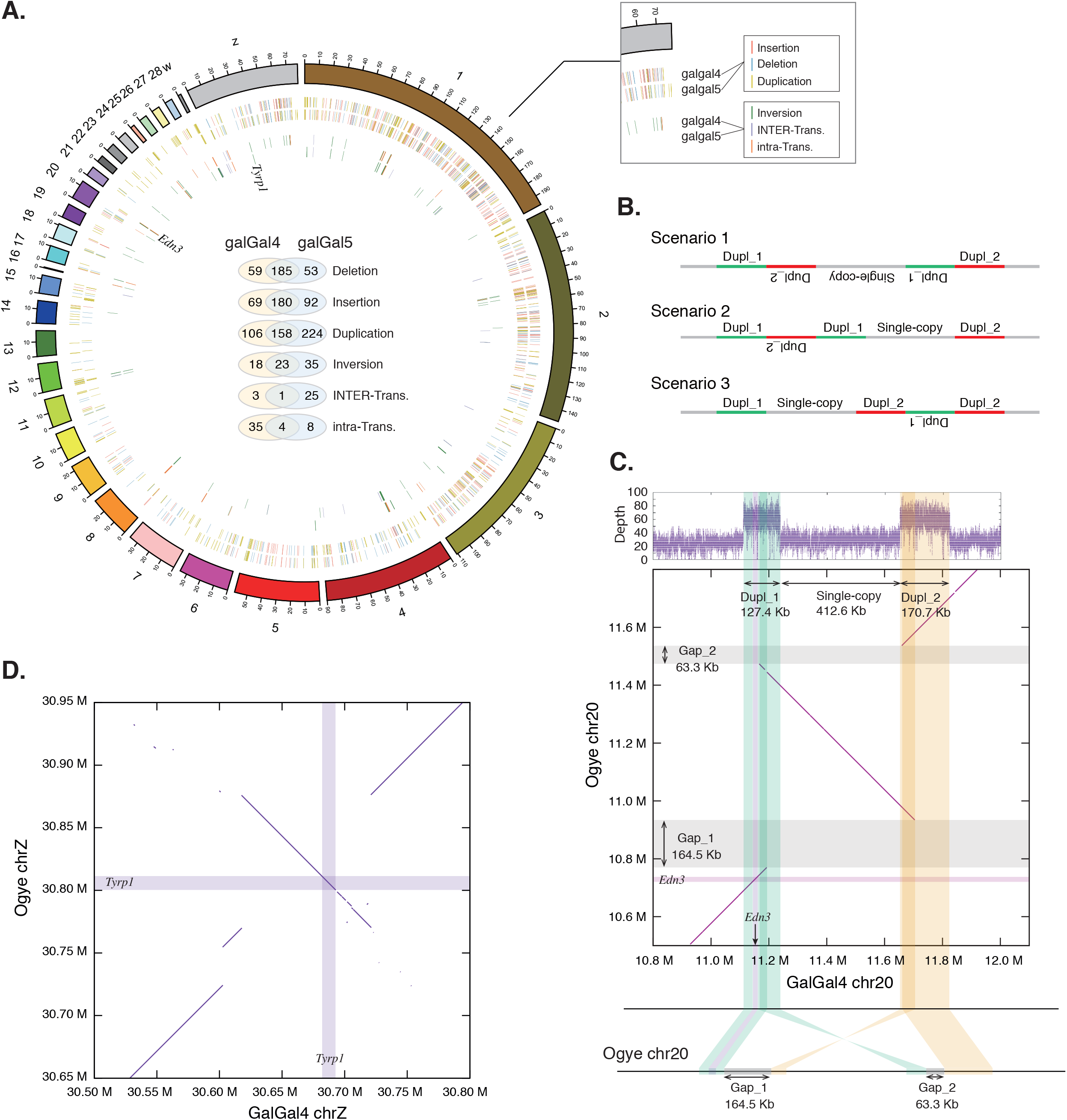
**A.** Structural variation (SV) map of the Ogye_1 genome compared with galGal4 and galGal5. Insertions (red), deletions (green), duplications (yellow), inversions (blue), inter-chromosomal translocations (gray; inter-trans), and intra-chromosomal translocations (orange; intra-trans) are shown. Variations in common between the genomes are shown in the middle with Venn diagrams; **B.** Three possible scenarios that could have led to SV (inverted duplication) of the Fibromelanosis (*FM*) locus in the genomes of hyperpigmented chicken breeds; **C.** Copy gain of the *FM* locus, which includes the *end3* gene (indicated by the thin purple-shaded boxes), was identified on chromosome 20. The green- and yellow-shaded boxes indicate duplicated regions (Dupl_1 and Dupl_2, respectively) and the gray-shaded boxes indicate gaps (Gap_1 and Gap_2). The sizes of Gap_1 and Gap_2 were estimated to be 164.5 Kbp and 63.3 Kbp, respectively; **D.** Inversion of a genomic region on chromosome Z that includes *tyrp1.* The purple-shaded boxes indicate the *tyrp1* locus.

Although the Fibromelanosis (*FM*) locus, which contains the hyperpigmentation-related *edn3* gene, is known to be duplicated in the genomes of certain hyperpigmented chicken breeds, such as *Silkie* and *Ayam cemani* [3, 6], the exact structure of the duplicated *FM* locus in such breeds has not been completely resolved due to its large size (~1Mbp). A previous study suggested that the inverted duplication of the *FM* locus could be explained by three possible mechanistic scenarios (**Figure 2B**) [3]. To understand more about the mechanism of *FM* locus SV in the Ogye_1 genome, we compared it to the same locus in the galGal4 genome. Aligning paired-end reads of the Ogye_1 genome to the galGal4 genome, we found higher read depth at the *FM* locus in *YO*, indicating a gain of copy number at the locus (**Figure 2C** top). Also, we have identified same mate-pair information using paired-end and mate-pair reads (see Supplementary **Figure S5**). The intervening region between the two duplicated regions was estimated to be 412.6 Kbp in length in Ogye_1. Regarding possible mechanistic scenarios, mate-pair reads (3-10 Kbp, and FOSMID) mapped to the locus supported all three suggested scenarios, but an alignment of chromosome 20 from Ogye_1 and galGal4 showed that the intervening regions, including inner-partial regions in both duplicated regions, were inverted at the same time, which supports the first mechanistic scenario in **Figure 2B**. Given the resulting alignments, the *FM* locus of the Ogye_1 genome was updated according to the first scenario. The size of Gap_1 and Gap_2 were estimated at 164.5 Kbp and 63.3 Kbp, respectively.

Additionally, a large inversion was detected near a locus including the *tyrp1* gene (**Figure 2D**), which is known to be related to melanogenesis [29, 30]. However, when resequencing data from *White Leghorn* (white skin and plumage), *Korean Black* (white skin and black plumage), and white *Silkie* (black skin and white plumage) were compared to the galGal4 or 5 genome assemblies, the same break points related to the inversion were detected (see supplementary **Figure S6** and **Figure S7**), suggesting that the inversion including *tyrp1* is not specifically related to skin hyperpigmentation.

### Annotations

#### Repeats

Repeat elements in the Ogye_1 and other genomes were predicted by a reference-guided approach, RepeatMasker [31], which utilizes Repbase libraries [32]. In the Ogye_1 genome, 205,684 retro-transposable elements (7.65%), including LINEs (6.41%), SINEs (0.04%) and LTR elements (1.20%), 27,348 DNA transposons (0.94%), 7,721 simple repeats (0.12%), and 298 low-complexity repeats (0.01%) were annotated (**Figure 3**, Supplementary **Table S2** and **Table S3**). It turns out that the composition of repeats in the Ogye_1 genome is similar to that in other avian genomes (**Figure 3**). Compared with other avian genomes, the Ogye_1 genome is more similar to galGal4 and 5 in terms of repeat composition including transposable elements (TEs) except for the fractions of simple repeats (0.12% for Ogye_1, 1.12% for galGal4 and 1.24% for galGal5), low-complexity (0.01% for Ogye_1, 0.24% for galGal4 and 0.25% for galGal5) and satellite DNA repeats (0.01% for Ogye_1, 0.20% for galGal4 and 0.22% for galGal5). The density of TEs across all chromosomes was depicted in **Figure 4A**.

**Figure 3.**
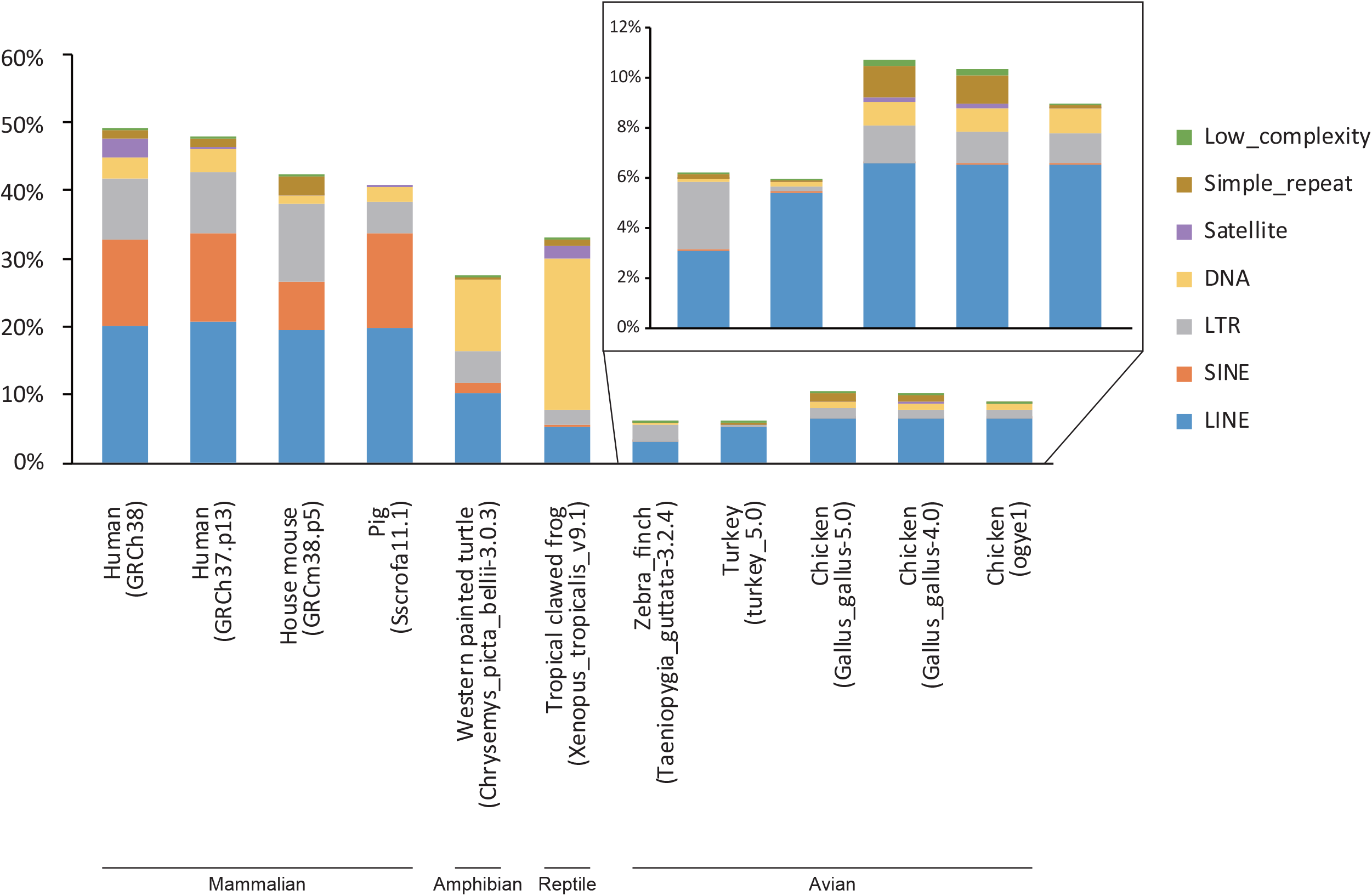
Composition of repeat elements in different assemblies of avian, reptile, and mammalian genomes. The repeats in unplaced scaffolds were not considered.

**Figure 4.**
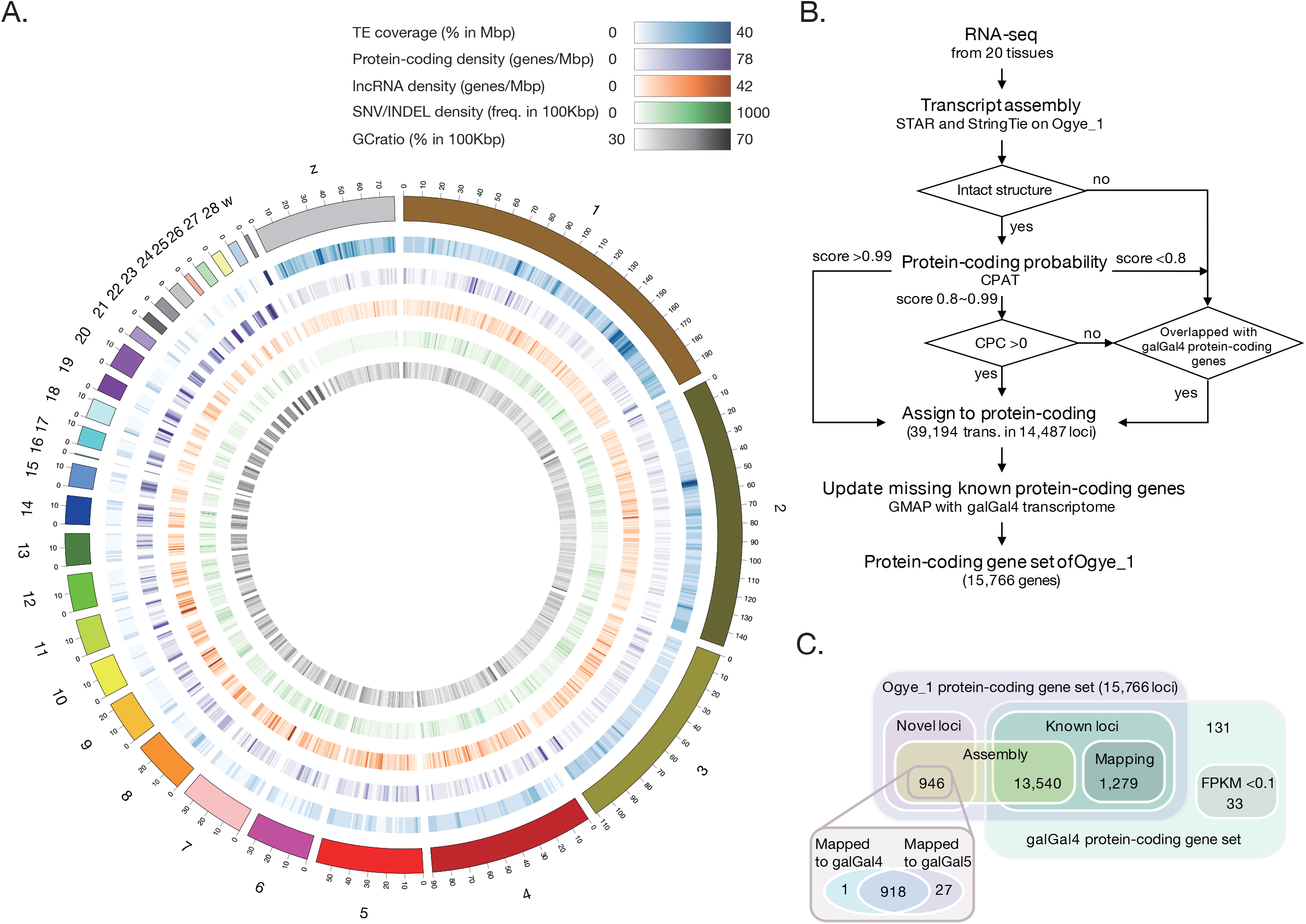
**A.** Gene (protein-coding and lncRNA) annotation maps of the Ogye_1 genome with TE, SNV/INDEL, and GC ratio landscapes are shown in a Circos plot. Note that TE and SNV denote transposable element and single nucleotide variation, respectively; **B.** A schematic flow of our protein-coding gene annotation pipeline and a Venn diagram showing the number of protein-coding genes in the Ogye_1 genome. **C.** We have annotated 946 novel genes, and found 13,541 known genes by transcript assembly. 1,279 known genes were annotated by mapping using GMAP. 164 annotated genes were not included in our *YO* protein-coding gene set, among which 33 were not expressed (<0.1 FPKM). All of the 946 newly annotated genes are mapped to the galGal4 or galGal5 genomes.

#### SNPs/INDELs

To annotate SNPs and INDELs in the Ogye_1 genome, we mapped all paired-end libraries (SRR6189087-SRR6189094) to the Ogye_1 genome using BWA-MEM, and performed a series of postprocesses including deduplication by Picard modules [33]. As a result, we have annotated 895,988 SNPs and 82,392 insertions/deletions (INDELs) across the genome using Genome Analysis Toolkit (GATK) modules. In performing GATK, we used HaplotypeCaller, combineGVCF, GenotypeGVCFs and VariantFiltration (with options `“QD < 2.0 || FS > 200.0 || ReadPosRankSum < -2 0.0”`) [22]. The densities of SNPs and INDELs across all chromosomes are depicted in **Figure 4A**.

#### Protein-coding genes

By analyzing large-scale of RNA-seq data from twenty different tissues through our protein-coding gene annotation pipeline (**Figure 4B**), 15,766 protein-coding genes were annotated in the Ogye_1 genome (**Figure 4C**), including 946 novel genes and 14,820 known genes. 164 protein-coding genes annotated in galGal4 were missing from the Ogye_1 assembly. The density of protein-coding genes across all chromosomes was depicted in **Figure 4A**.

To sensitively annotate protein-coding genes, all paired-end RNA-seq data were mapped on the Ogye_1 genome by STAR [34] for each tissue and the mapping results were then assembled into potential transcripts using StringTie [35]. Assembled transcripts from each sample were merged using StringTie and the resulting transcriptome was subjected to the prediction of coding DNA sequences (CDSs) using TransDecoder [36]. For high-confidence prediction, transcripts with intact gene structures (5’UTR, CDS, and 3’UTR) were selected. To verify the coding potential, the candidate sequences were examined using CPAT [37] and CPC [38]. Candidates with a high CPAT score (>0.99) were directly assigned to be protein-coding genes, and those with an intermediate score (0.8-0.99) were re-examined to determine whether the CPC score is >0. Candidates with low coding potential or that were partially annotated were examined to determine if their loci overlapped with annotated protein-coding genes from galGal4 (ENSEMBL cDNA release 85). Overlapping genes were added to the set of Ogye_1 protein-coding genes. Finally, 164 genes were not mapped to the Ogye_1 genome by GMAP, 131 of which were confirmed to be expressed in *YO* (≥ 0.1 FPKM) using all paired-end *YO* RNA-seq data. However, expression of the remaining 33 genes was not confirmed, suggesting that they are not expressed in *YO* (< 0.1 FPKM) or have been lost from the Ogye_1 genome. The missing genes are listed in **Table S4**. In contrast, the 946 newly annotated Ogye_1 genes appeared to be mapped to the galGal4 or galGal5 genomes (**Figure 4C**).

#### lncRNAs

In total, 6,900 *YO* lncRNA genes, including 5,610 novel loci and 1,290 known loci, were confidently annotated from RNA-seq data using our lncRNA annotation pipeline, adopted from our previous study [39] (**Figure 5A**).

**Figure 5.**
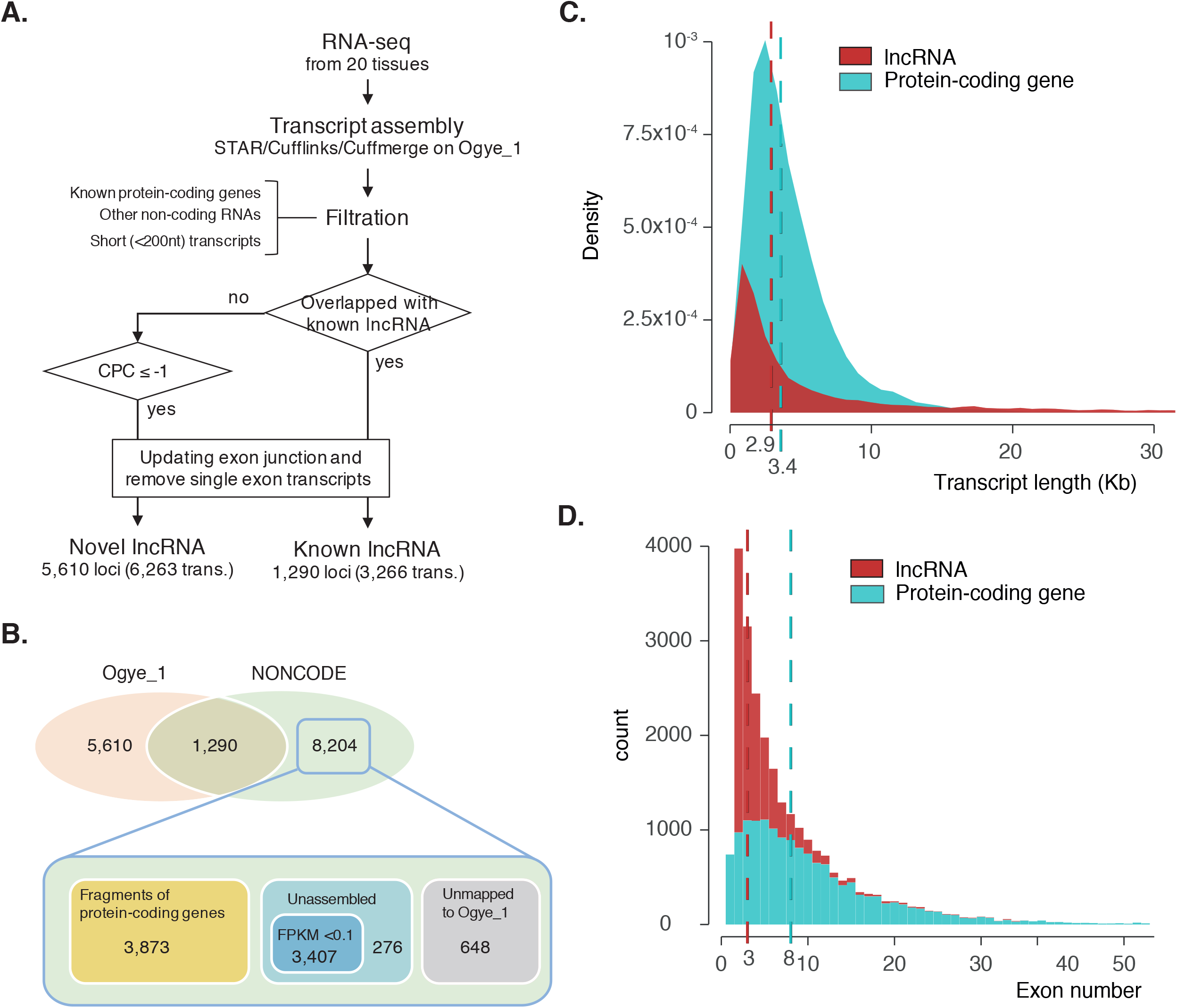
**A.** A computational pipeline for lncRNA annotations. **B.** The number of Ogye_1 and galGal4 lncRNAs are shown in the Venn diagram. About 51% (3,873) of non-overlapping galGal4 lncRNAs are predicted to be fragments of protein-coding genes. 45% (3,407) were not expressed in any tissue. Only 4% (276) of the lncRNAs are actually missing from the set of Ogye lncRNAs (bottom). **C.** Distribution of transcript length (red for lncRNAs and cyan for protein-coding genes). The vertical dotted lines indicate the median length. **D.** Distribution of exon number per transcript. Otherwise, as in **C.**

To annotated and profile lncRNA genes, pooled single- and paired-end RNA-seq reads of each tissue were mapped to the Ogye_1 genome (PRJNA412424) using STAR [34], and subjected to transcriptome assembly using Cufflinks [44], leading to the construction of transcriptome maps for twenty tissues. The resulting maps were combined using Cuffmerge and, in total, 206,084 transcripts from 103,405 loci were reconstructed in the Ogye genome. We removed other biotypes of RNAs (the sequences of mRNAs, tRNAs, rRNAs, snoRNAs, miRNAs, and other small non-coding RNAs downloaded from ENSEMBL biomart) and short transcripts (less than 200nt in length). 54,760 lncRNA candidate loci (60,257 transcripts) were retained, and which were compared with a chicken lncRNA annotation of NONCODE (v2016) [45]. Of the candidates, 2,094 loci (5,215 transcripts) overlapped with previously annotated chicken lncRNAs. 52,666 non-overlapping loci (55,042 transcripts) were further examined to determine whether they had coding potential using CPC score [38]. Those with a score greater than -1 were filtered out, and the remainder (14,108 novel lncRNA candidate loci without coding potential) were subjected to the next step. Because many candidates still appeared to be fragmented, those with a single exon but with neighboring candidates within 36,873bp, which is the intron length of the 99th percentile, were re-examined using both exon-junction reads consistently presented over twenty tissues and the maximum entropy score [46], as done in our previous study [39]. If there were at least two junction reads spanning two neighboring transcripts or if the entropy score was greater than 4.66 in the interspace, two candidates were reconnected, and those with a single exon were discarded. In the final version, 9,529 transcripts from 6,900 lncRNA loci (5,610 novel and 1,290 known) were annotated as lncRNAs (see **Figure 5B**), which included 6,170 (89.40 %) intergenic lncRNAs and 730 (10.57 %) antisense ncRNAs. Consistent with other species [40-43], the median gene length and the median exon number of Ogye lncRNAs were less than those of protein-coding genes (**Figure 5C** and **D**).

Although 13,540 of 15,766 protein-coding genes (92%) were redetected by our transcriptome assembly and protein-coding gene annotations (see **Figure 4C**), 8,204 of NONCODE lncRNAs were missed in our Ogye lncRNA annotations (**Figure 5B**), the majority of which were either fragments of protein-coding genes or not expressed in all twenty Ogye tissues (**Figure 5B**). Only 276 were missed in our transcriptome assembly, and 648 were missed in our draft genome.

### Coding and non-coding transcriptome maps

Using paired-end *YO* RNA-seq data, the expression levels of protein-coding and lncRNA genes were calculated across twenty tissues (**Figure 6A**), which were dynamically changed. The profiled transcriptomes included 1,814 protein-coding genes and 1,226 lncRNA genes, expressed ≥ 10 FPKM in only one tissue as well as 1,559 and 351 expressed ≥ 10 FPKM in all tissues. The Ogye black tissues (fascia, comb, skin, and shank) expressed 6,702 protein-coding and 3,291 lncRNA genes ≥ 10 FPKM, the majority of which appeared to be expressed in tissue-specific manner (**Figure 6B**). For instance, two neighboring genes, *krt9* and *lnc-lama2-1* are highly expressed in comb and shank, and (**Figure 6C** and **D**).

**Figure 6.**
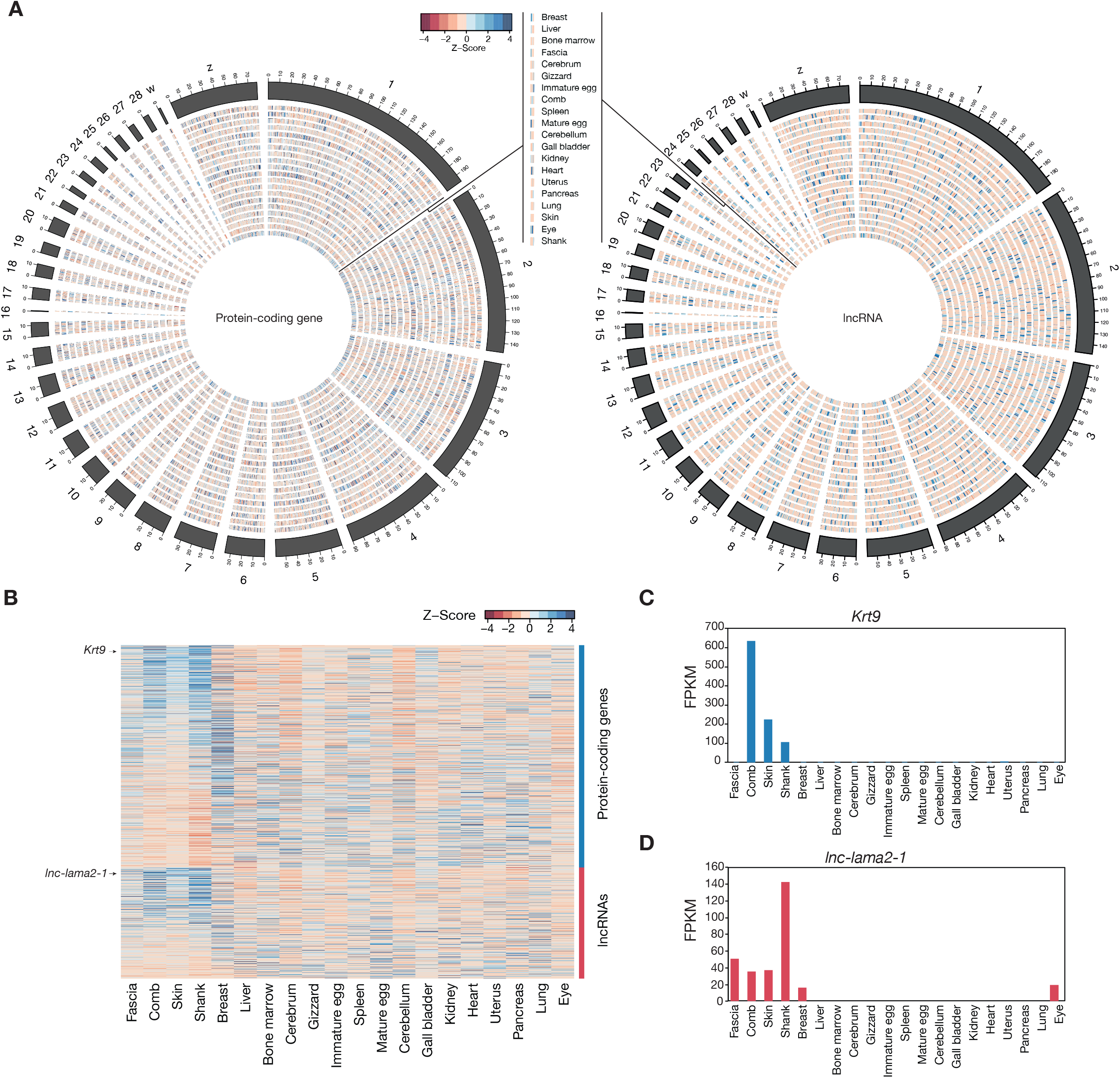
**A.** The circus plot illustrates the expression levels of protein-coding genes (left) and lncRNA (right) across twenty tissues. The expression levels are indicated with a color-coded z-score, described in the key; **B.** The expression patterns of black tissues-specific genes. Expression levels are indicated with a color-coded Z-score (red for low and blue for high expression) as shown in the key; **C.** Expression levels of *krt9* across twenty tissues; **D.** Expression levels of *lnc-lama2-1* across twenty tissues.

As lncRNAs tend to be specifically expressed in a tissue or in related tissues, they could be better factors for defining genomic characteristics of tissues than protein-coding genes. To prove this idea, principle component analyses (PCA) were performed with tissue-specific lncRNAs and protein-coding genes using reshape2 R package (**Figure 7**) [47]. As expected, the 1st, 2nd, and 3rd PCs of lncRNAs enabled us to predict the majority of variances, and better discerned distantly-related tissues and functionally and histologically-related tissues (i.e., black tissues and brain tissues) (**Figure 7B**) than those of protein-coding genes (**Figure 7A**).

**Figure 7.**
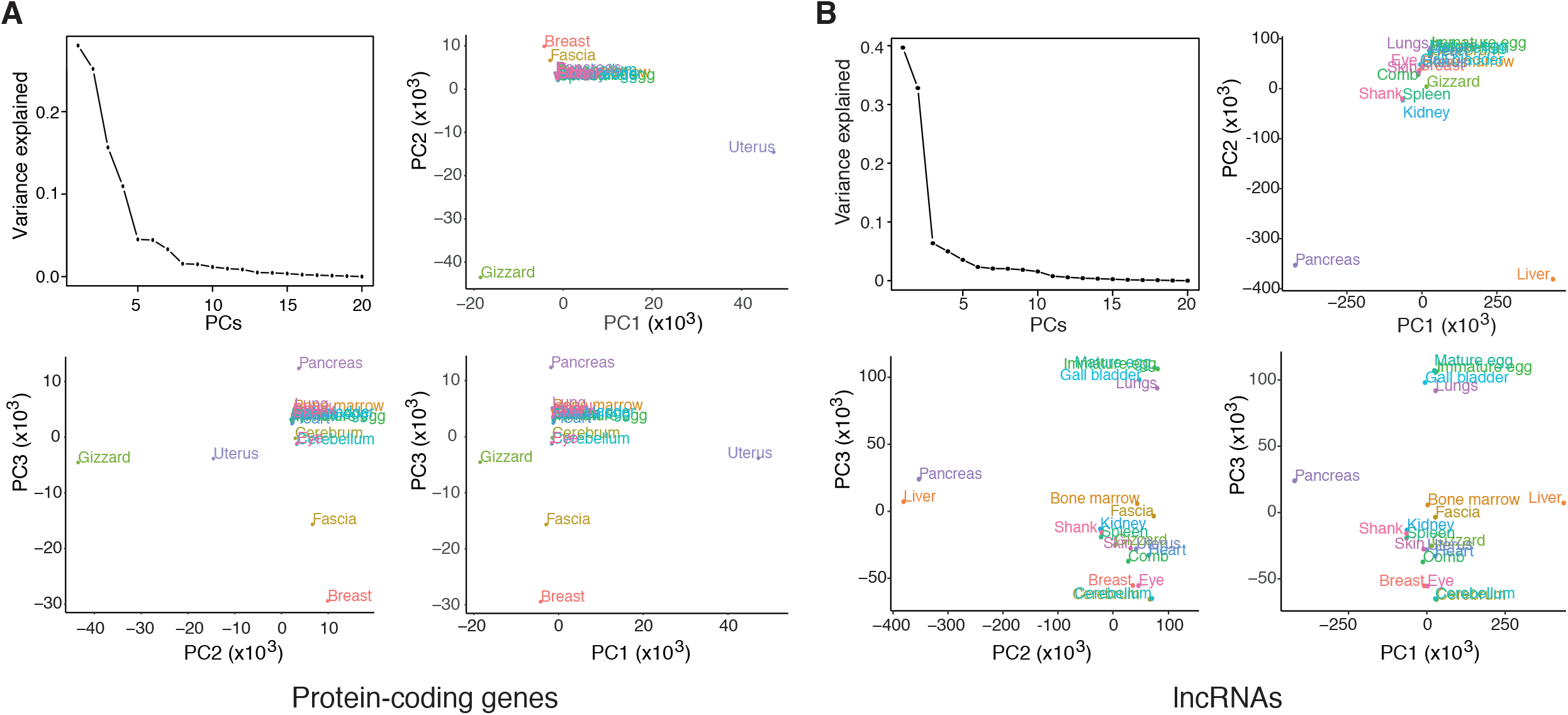
**A.** Principal component analysis (PCA) using tissue-specific lncRNAs. PCs explaining the variances are indicated with the amount of the contribution in the left-top plot. PCA plots with PC1, PC2, and PC3 were demonstrated in a pairwise manner. Each tissue is indicated on the PCA plot with a specific color; **B.** PCA using tissue-specific protein-coding genes. Otherwise, as in **A.**

### DNA methylation maps

RRBS reads were mapped to Ogye draft genome (**Table 3**), and calculated DNA methylation signals (C to T changes in CpGs) across chromosomes using Bismark in each sample [48]. The DNA methylation landscape in protein-coding and lncRNA genes were shown in **Figure 8A**. Based on CpG methylation pattern in the promoter of genes, hierarchical clustering was performed using rsgcc R package, and clusters including adjacent or functionally similar tissues, such as cerebrum and cerebellum, immature and mature eggs, or comb and skin were identified (**Figure 8B**). Of all CpG sites in genomes, 31~65% were methylated across tissues while only 19~43% were methylated in the promoter (**Table 5**), indicating hypomethylation status in the promoters of expressed genes. Exceptionally, the methylation levels of spleen (40.28% in all genomic regions; 24.92% in promoter region) and liver (30.82% in all genomic region; 18.68% in promoter regions) displayed much less methylation signal, compared to those of others. In fact, examining the averaged methylation landscapes over protein-coding and lncRNA loci showed that the methylation levels in gene body regions were much higher than those in 2 Kbp upstream regions from transcript start sites (TSS) (**Figure 8C** and D). To confirm that CpG methylation represses the expression, 280 protein-coding and 392 lncRNA genes of *Max*[*FPKM*] ≥ 10 with highly tissue-specific expression pattern were selected. The methylation level of highly expressed genes was much lower than others (**Figure 8E** and **F**).

**Table 5.**
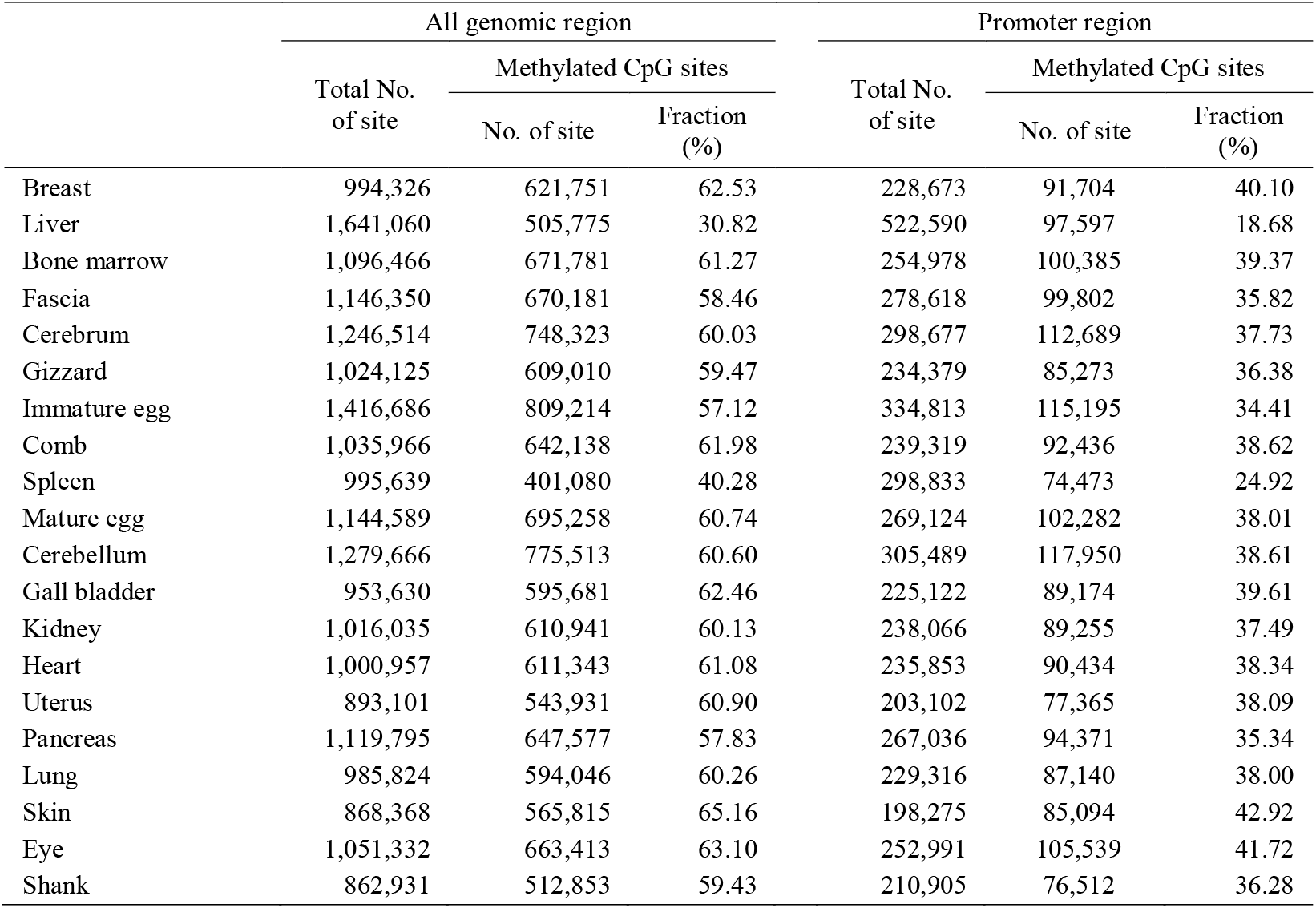
Summary of methylated CpGs across twenty tissues.

**Figure 8.**
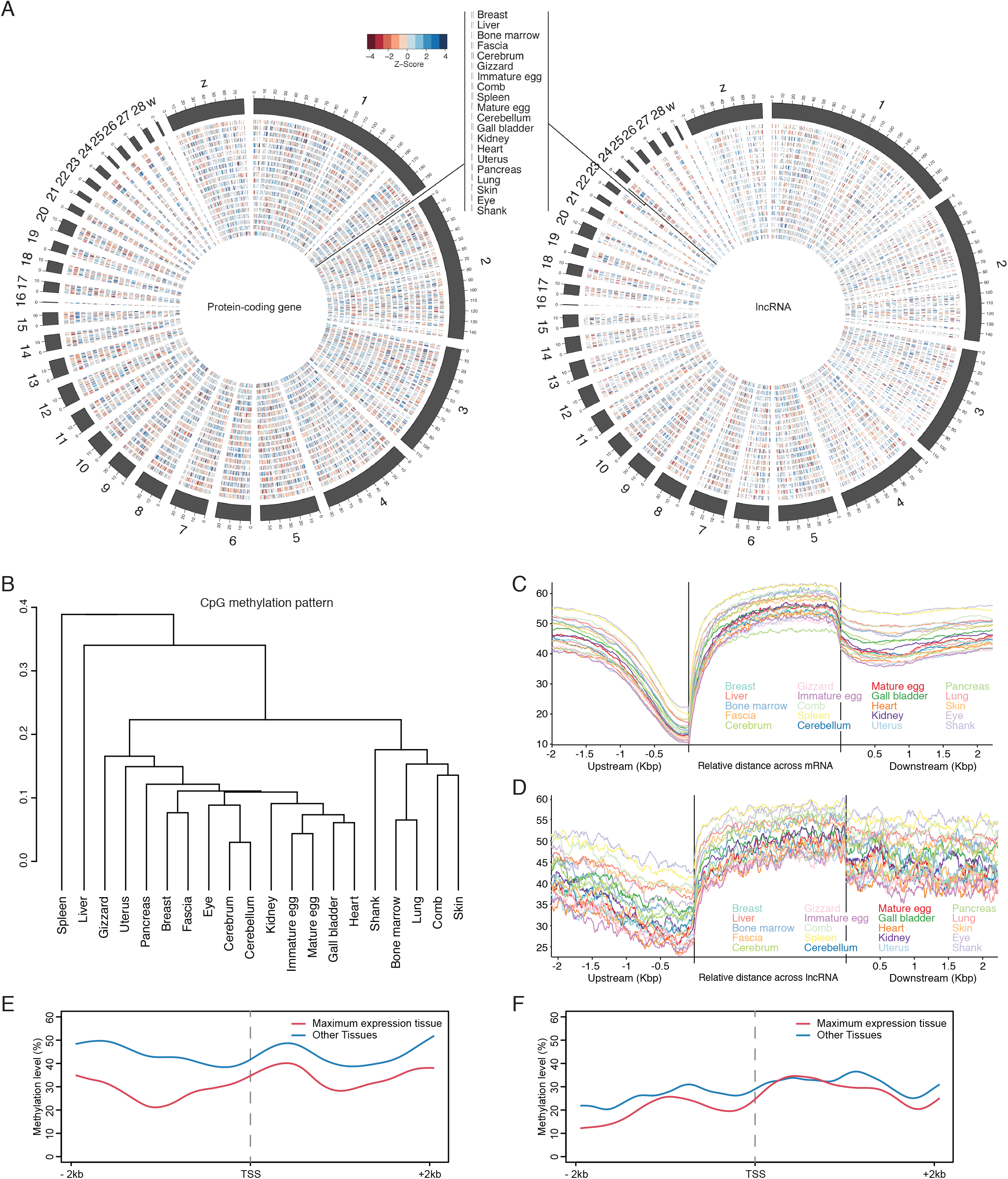
**A.** The circus plot illustrates the CpG methylation levels in the promoters of protein-coding genes (left) and lncRNA (right) across twenty tissues. The methylation levels are indicated with a color-coded z-score, described in the key; **B.** Hierarchical clustering using Pearson correlation of DNA methylation patterns between tissues; **C.** Average DNA methylation landscape across gene bodies of protein-coding genes and their flanking regions; **D.** Average DNA methylation landscape across gene bodies of lncRNAs and their flanking regions; **E.** Average DNA methylation level of the protein-coding gene in tissues where the gene is highest expressed and lowest expressed, respectively. **F.** Average DNA methylation level of the lncRNA gene in tissues where the gene is highest expressed and lowest expressed, respectively. The methylation level is indicated with a color-coded z-score, described in the key.

To correlate the expression of genes with their methylation levels, only tissues in which a certain position had sufficient read coverage (at least five) were considered for measuring the correlation. The Spearman correlation coefficients between the expression and methylations levels were observed over twenty tissues (**Figure 9**). 454 protein-coding and 25 lncRNA genes displayed a negative correlation to methylation levels in promoter regions, while 157 protein-coding and 20 lncRNA genes have a positive correlation (box plots in **Figure 9**)

**Figure 9.**
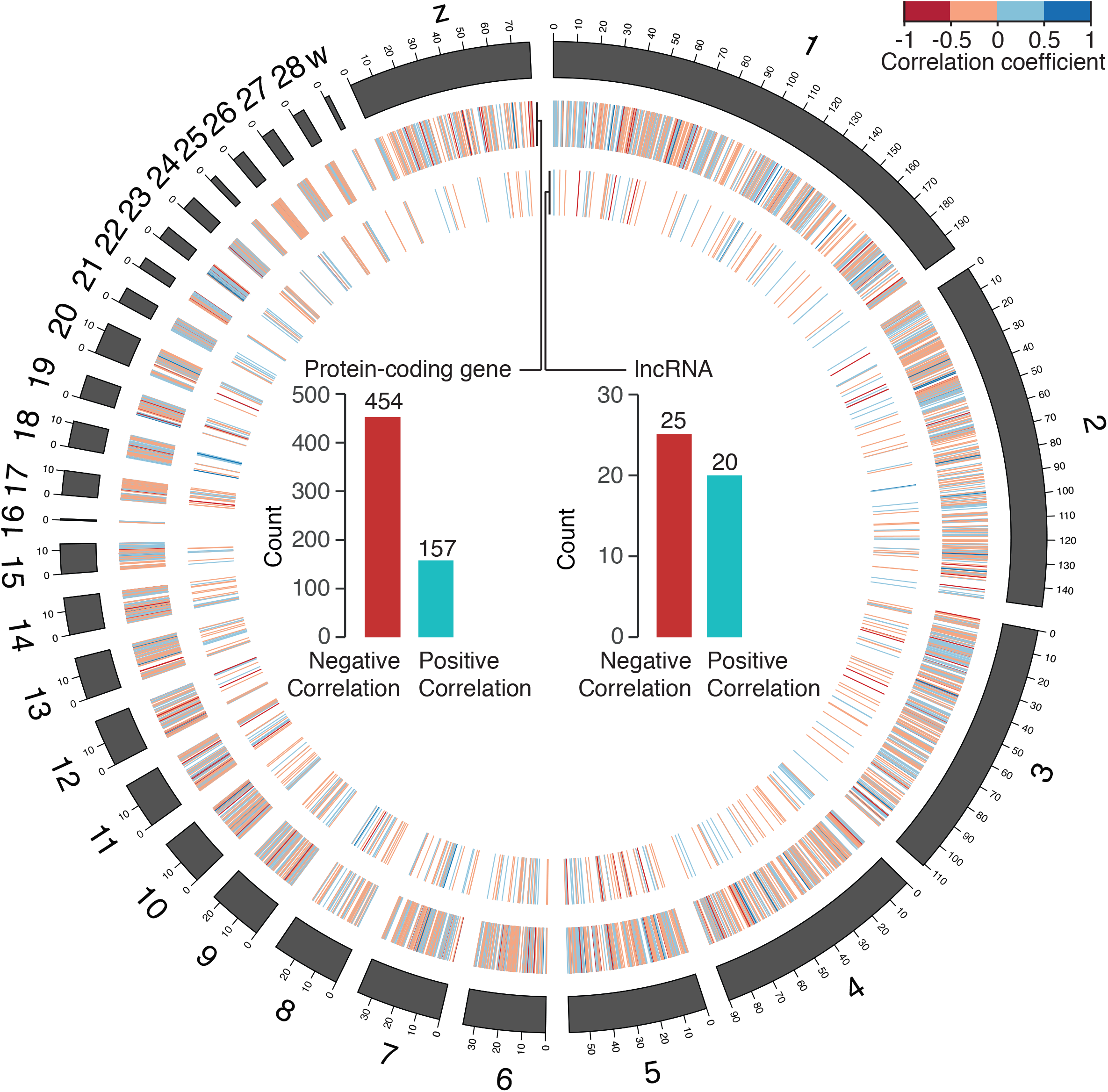
A circos plot illustrating the Spearman correlation coefficients between expression levels and methylation levels of genes across chromosomes (heatmaps). The bar chart indicates the count of genes (left for protein-coding genes and right for lncRNAs) with a significant negative (red) and positive (cyan) correlation between the methylation level in their promoters and their expression values.

### Discussions

In this work, the first draft genome from a Korean native chicken breed, *YO*, was constructed with genomic variation, repeat, and protein-coding and non-coding gene maps. Compared with the chicken reference genome maps, many more novel coding and non-coding elements were identified from large-scale RNA-seq datasets across twenty different tissues. Although the Ogye_1 genome is comparable with galGal5 with respect to genome completeness evaluated by BUSCO, the Ogye_1 seems to lack simple and long repeats compared with galGal5, which was assembled from high-depth PacBio long-reads (50X) [10] that can capture simple and long repeats. Although we also used PacBio long reads, some simple and satellite repeats would be missed during assembly, because we used the PacBio data of shallow depth (11.5X) for scaffolding and gap filling. A similar tendency can be seen in the Golden-collared manakin genome (ASM171598v1) (**Figure 3**), which was also assembled in a hybrid manner using MaSuRCA assembler with high-depth Illumina short-reads and low-depth PacBio long-reads.

15,766 protein-coding 6,900 lncRNA genes were annotated from twenty tissues of *YO.* 946 novel protein-coding genes were identified while 164 genes of *Galllus gallus red junglefowl* were missed in our annotations. In the case of lncRNAs, only about 19% of previously annotated chicken lncRNAs were redetected, and the remainders were mostly not expressed in *YO* or were false annotations, suggesting that the current chicken lncRNA annotations should be carefully examined. Our Ogye lncRNAs resembled previously annotated lncRNAs in mammals in their genomic characteristics, including transcript length, exon number, and tissue-specific expression pattern, providing evidence for the accuracy of the new annotations. Hence, our lncRNA catalogue may help us to improve lncRNA annotations in the chicken reference genome.

Annotated genomic variations and comparative analysis of coding and non-coding genes will provide a resource for understanding genomic differences and evolution of *YO* as well as identifying functional elements related to its unique phenotypes, such as fibromelanosis. Additionally, such analyses will be useful for future genome-based animal genetics.

### Availability of data

All of our sequencing data and the genome sequence have been deposited in NCBI’s BIOPROJECT under the accession number PRJNA412424. The raw sequence data have been deposited in the Short Read Archive (SRA) under accession numbers SRR6189081-SRR6189098 (**Table 1**), SRX3223583-SRX3223622 (**Table 2**), and SRX3223667-SRX3223686 (**Table 3**).

### Additional files

The supplementary Figures and tables have been included in a supplementary file:

**Figure S1.** Assembly statistics of Ogye_1 gnome assembly at each step.

**Figure S2.** Filtration of noise and mis-assembly detection using Lastz

**Figure S3.** Hierarchical mapping information in the reference-assisted additional assembly pipeline.

**Figure S4.** Alignment of the Ogye_1 genome to galGal4/5 drawn by MUMmer.

**Figure S5.** Mate-pair information in *FM* locus.

**Figure S6.** Mate-pair information near tyrp1 in chrZ of galGal4 and galGal5.

**Figure S7.** Break points near *tyrp1* in galGal4 chrZ.

**Table S1.** Structural variations in the Ogye_1 genome

**Table S2.** Repeats in the Ogye_1 genome

**Table S3.** Repeat contents in different assemblies.

**Table S4.** 164 unmapped genes among galGal4 protein-coding genes.

## Acknowledgements

We thank all members of the BIG lab for helpful comments and discussions. This work was supported by the Cooperative Research Program for Agriculture Science and Technology Development (Project title: National Agricultural Genome Program, Project No. PJ01045301 and PJ01045303).

## Author’s Contributions

KTL, NSK, HHC, and JWN designed the study, KTL, YJD and CYC collected samples, DJL, HHC and KTL collected sequencing data, and JIS, KWN, NSK, JMK, HHC and JMN performed the analysis and developed the methodology. JIS, KWN and JMK wrote the manuscript.

